# Rns regulates nonpilus adhesin EtpA in human Enterotoxigenic *E.coli* (ETEC)

**DOI:** 10.1101/362178

**Authors:** T.P.Vipin Madhavan

**Affiliations:** School of Medical Science, Griffith University, Gold Coast Campus, Australia QLD4222

## Abstract

Rns, an araC family of transcriptional activator (AFTR) is known to regulate many of the known pili in human ETEC. Apart from pili, Rns is also known to regulate some nonpilus genes believed to have role in virulence. EtpA is a nonpilus adhesin, encoded with in *etpBAC* operon in ETEC genome. Using a combination of qRT-PCR and gel shift assay, we show that Rns binds to upstream of etpBAC operon and upregulates the expression of EtpA. This is the first report of Rns regulating a known virulence factor in ETEC.

## Introduction

AraC family of transcriptional activators (AFTRs) regulates a large number of genes, belonging to different metabolic pathways in different bacteria (Gallegos et al., 1997). For example, the AFTR, PerA, regulates more than 40 genes required for Bfp pilus assembly, a Type III secretion system and secreted virulence proteins (Porter et al., 2004). In *Vibrio cholerae*, ToxR controls the expression of virulence factors such as Tcp pili and cholera toxin as well as several other genes with potential or confirmed roles in virulence (Zhang et al., 2004, Zhang et al., 2003a, Zhang et al., 2003b). Similarly, the AFTR, RegA, from *Citrobacter rodentium* regulates the expression of more than 30 genes, including virulence genes encoding pilus and non-pilus adhesins, potential virulence factors and house-keeping genes (Yang et al., 2011, Yang et al., 2010, Hart et al., 2008). Therefore, in general, AFTRs controlling pilus expression in these pathogens also control the expression of large sets of non-pilus genes, including genes that encode other types of virulence factor.

Rns is an AFTR that was originally identified as a positively regulator of the expression of CS1 and CS2 pili in ETEC (Caron et al., 1989). Subsequent studies showed that Rns could regulate almost half of the known pilus systems in ETEC (Basturea et al., 2008), as well as four ETEC genes *cexE, yiiS, nlpA*, and *aslA* that do not encode pili (Pilonieta et al., 2007, Bodero et al., 2007, Munson et al., 2002). By analogy to Rns homologues in other bacteria, we hypothesized that Rns also controls the expression of many virulence factors that are not reported yet. Using a transcriptomic comparison between an ETEC WT strain and a single crossover mutant of Rns, we showed that Rns is upregulates the expression of non-pilus adhesin, EtpA and its partner outer membrane protein, EtpB. Further bioinformatic and gel shift assays showed that Rns binds to upstream of etpBAC operon.

## Materials and Methods

### Bacterial strains, plasmids and culture conditions

Cultures were grown aerated in Luria-Bertani (LB) broth or agar (Scott, 1974), or 2YT broth (Sambrook and Russell, 2001), or CFA agar or broth (Evans et al., 1980) at 37° C with antibiotic selection where appropriate. The *E. coli K*-12 strains DH5α (Hanahan, 1983) and DH5α/F’-*lacI^q^* (New England Biolabs) were used as general hosts for cloning. *E. coli* strain BL21(DE3) Rosetta (Novagen) was used for high-level expression under the T7 promoter. Further details of the strains are given in Table 1. Plasmids used or constructed in this study are listed in Table 2.

**Table 1:** List of bacterial strains

| Strain | Genotype/Characteristics | Reference/<br>Source |
| --- | --- | --- |
| *E. coli* DH5α/F'-*lacI<sup>q</sup>* | F'[*lacI<sup>q</sup>*] *endA1 glnV44 thi-1 recA1 relA1 gyrA96 deoR nupG Φ80dlacZΔM15 $\Delta$(lacZYA-argF)U169, hsdR17(r<sub>K</sub><sup>-</sup> m<sub>K</sub><sup>+</sup>), λ<sup>-</sup>* | Novagen |
| *E. coli* BL21 DE3 Rosetta | F<sup>-</sup> *ompT gal dcm lon hsdS<sub>B</sub>(r<sub>B</sub><sup>-</sup> m<sub>B</sub><sup>-</sup>) $\lambda$(DE3 [*lacI lacUV5-T7* gene 1 *ind1 sam7 nin5*]) pSRARE (*cat*)* | Novagen |
| *E. coli* B2C | O6:H16:CFA/II strain of enterotoxigenic *E. coli* | (DuPont et al., 1971) |
| *E. coli* B2C Rif<sup>r</sup> | Spontaneous rifampicin resistant mutant of B2C | H. Sakellaris |
| *E. coli* B2C Rif<sup>r</sup> *rns* | ETEC B2C Rif<sup>r</sup>::pCactus-*mob-rns*'; single cross-over homologous recombination of pCactus-*mob-rns*' into chromosome inactivates *rns* gene. | H. Sakellaris |

**Table 2:** List of plasmids used in the study.

| Plasmid | Backbone<br>Vector | Relevant Characteristics | Reference/<br>Source |
| --- | --- | --- | --- |
| pMAL-c2X |  | P<sub>tac</sub>-*malE-lacZ'* | New England Biolabs |
| pBBR1MCS-2 | pBBR1MCS |  | (Kovach et al., 1995) |

### Construction of single cross-over mutant of rns

An insertional mutation of *rns* was constructed by selecting for a single cross-over homologous recombination of the suicide plasmid pCactus-*mob-rns’* into the chromosome of B2C Rif^r^. In order to confirm the mutation, Genomic DNA of putative ETEC B2C *rns* mutants (ETEC B2C Rif/pCactus-*mob*-*rns’*) and ETEC B2C Rif^r^ was extracted using Fermentas GeneJET genomic DNA purification kit (Thermo Fisher Scientific, Australia), according to the manufacturers’ protocol. Primers homologues to 5’upstream and 3’ downstream of *rns* was used to amplify the *rns* loci. Insertion of pCactus-*mob*-*rns’* was confirmed using agarose gel electrophoresis based on the difference in size of amplicons of wild type and *rns* mutant. Abrogation of rns expression was confirmed by qRT-PCR (data not shown).

### RNA extraction and purification

Bacterial strains ETEC B2C WT/pBBR1MCS and ETEC B2C Rif/pCactus-mob-rns’ were grown in 3mL of colonisation factor antigen (CFA media) overnight at 37°C in a shaking incubator to make a primary inoculum. To 20ml of CFA broth, primary inoculum was inoculated at 1 % (v/v) concentration, incubated until an OD_600_ of 0.4 was reached. Two volumes of bacterial RNA protection reagent (Qiagen RNAprotect Bacteria Reagent) was added and the culture was centrifuged at 14,000 x *g*. The bacterial cell pellet was frozen at −80°C. Bacterial RNA was extracted from the frozen pellets using the Promega SV Total RNA Purification kit. Genomic DNA was removed using TURBO™ DNase (Thermo Fisher Scientific) following the manufacturers guidelines. Final RNA samples were quantified using. NanoDrop (NanoDrop products, USA).

### qRT-PCR

cDNA was synthesised using M-MuLV Reverse Transcriptase (NEB, Australia) following the manufacturers guidelines. Reaction mix for qRT-PCR was prepared using SensiFAST SYBR No-ROX Kit (Bioline) and quantified using the LightCycler® 480 System (Roche) following the manufacture’s guidelines. The internal control for all qRT-PCR reactions was *gyrB*. The relative fold difference in cDNA from each sample was calculated using the 2^-ΔΔC_T_^ method as previously described (Livak and Schmittgen, 2001). Graphs were plotted using GraphPad Prism (version 6). Details of the primers used in this study are given in table 3.

### Construction and purification of MBP-Rns fusion protein

The *rns* gene was PCR amplified from ETEC B2C Rif^r^ genomic DNA using primers HSP279. The amplicon was purified, restricted with *Xba*I and *Hin*dIII and ligated with similarly restricted pMAL-C2X. Confirmed clone was transformed into BL21(DE3) Rosetta. Expression and purification was conducted as described previously (Munson and Scott, 1999).

### Electrophoretic mobility shift assay (EMSA)

Electrophoretic mobility shift assays (EMSA) were conducted as described previously (Munson and Scott, 1999) using an EMSA kit (Life Technologies), following manufacturers guidelines. Electrophoretic separation was conducted from 4 to 8 hours depending on the size of DNA fragment.

## Results

### Rns upregulates the expression of *etpA* and *etpB* in etpBAC operon

As pilus expression regulating AFTRs in a number of Gram-negative pathogens also known to regulate the transcription of non-pilus virulence factors, we tested the hypothesis that Rns regulates the transcription of a variety of known, non-pilus virulence factors in ETEC, using a directed approach with qRT-PCR. In this regard two virulence genes of etpBAC operon *etpA* and *etpB* were tested for Rns-regulation by qRT-PCR. The *etpA* and *etpB* genes encode a non-pilus adhesin and its transporter, respectively. The transcription of these genes was measured in B2C Rif^r^/pBBR1MCS-2 (wild-type), B2C Rif^r^ *rns/*pBBR1MCS-2 (*rns* mutant) and B2C Rif^r^ *rns/*pBBR1MCS-2-*rns* (complemented mutant). A comparison of the transcription of each gene was made in the *rns* mutant and wild-type strain, as well as in the *rns* mutant and complemented mutant strain. Transcription of *etpA*, and *etpB* was higher in the wild type and complemented mutant strains than it was in the *rns* mutant (Fig 1), indicating that Rns positively regulates the expression of these virulence genes. This was surprising as there have been no previous reports that these well-studied virulence genes are regulated by Rns. qRT-PCR measurement of the transcription of *etpA* and *etpB* in wild-type (WT) and complemented *rns* mutant (C) strains, relative to the *rns* mutant of B2C. The results indicate *that etpA,* and *etpB* are positively regulated by Rns.

**Fig 1:**
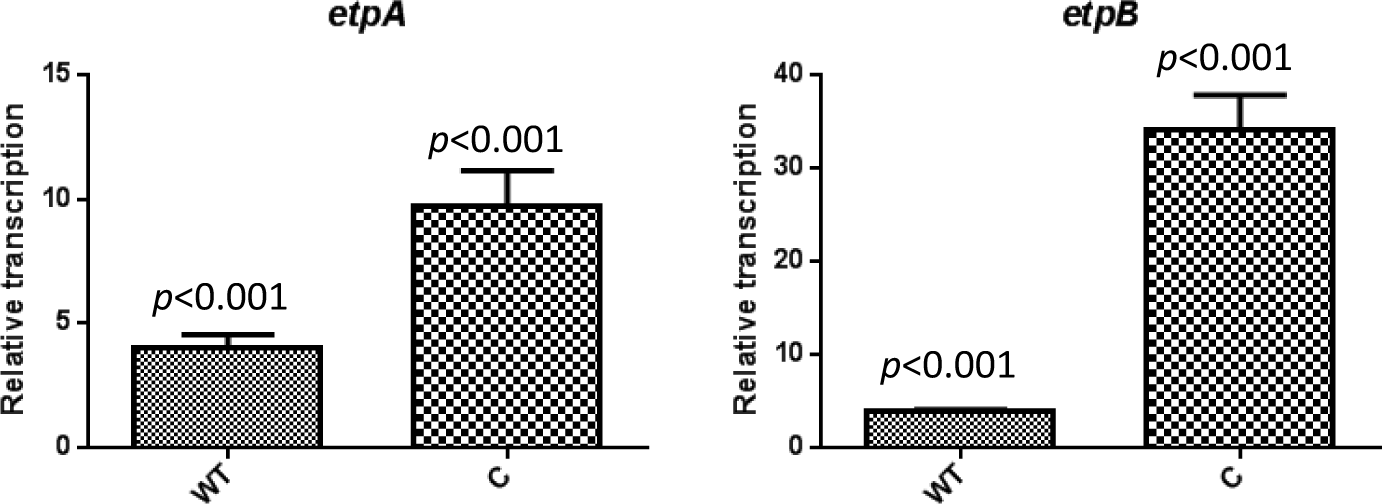
Rns-regulation of *etpA* and *etpB* genes in B2C.

### Rns binds to a DNA sequence upstream of the etpBAC operon

In order to study whether Rns regulates etpA and etpB genes directly, upstream sequences of etpBAC operon was analysed with searching algorithm, Fuzznuc (Rice et al., 2000), for the presence the Rns binding motif (Munson and Scott, 2000). The analysis revealed a putative Rns binding site 148bps upstream of *etpBAC* operon.

An alignment of the putative Rns-binding sequence with other Rns-binding sequences is shown in Fig 2. To test whether the putative Rns binding site was biologically significant, an electrophoretic mobility shift assay (See Methods) was conducted on a DNA fragment encompassing the sequence. A *malE-rns* fusion construct was cloned and MBP-Rns fusion protein was expressed and purified (See Methods) for use in EMSA. A test DNA fragment containing the putative Rns-binding site was PCR-amplified with the primers HSP411 and HSP412, and a negative control DNA sequence was generated by PCR-amplifying an internal fragment of *gyrB* with the primers HSP298 and HSP299. Reactions of MBP-Rns and DNA fragments were subjected to electrophoresis and DNA was visualized with SYBR Green staining (Fig 3). Reaction of the test DNA with MBP-Rns led to the formation of two high molecular weight complexes (Fig 3, lanes 2-4), that were absent from the negative control reaction containing only DNA (lane 1). As expected, increasing concentrations of MBP-Rns were accompanied by increasing concentrations of the higher molecular weight complexes and decreasing concentrations of uncomplexed DNA. The inability of MBP-Rns to cause a shift in the mobility of the negative control *gyrB* DNA fragment (Fig 3, lanes 5 and 6) indicated that the binding of MBP-Rns to the sequence upstream of the *etpBAC* operon was specific.

**Fig 2:**
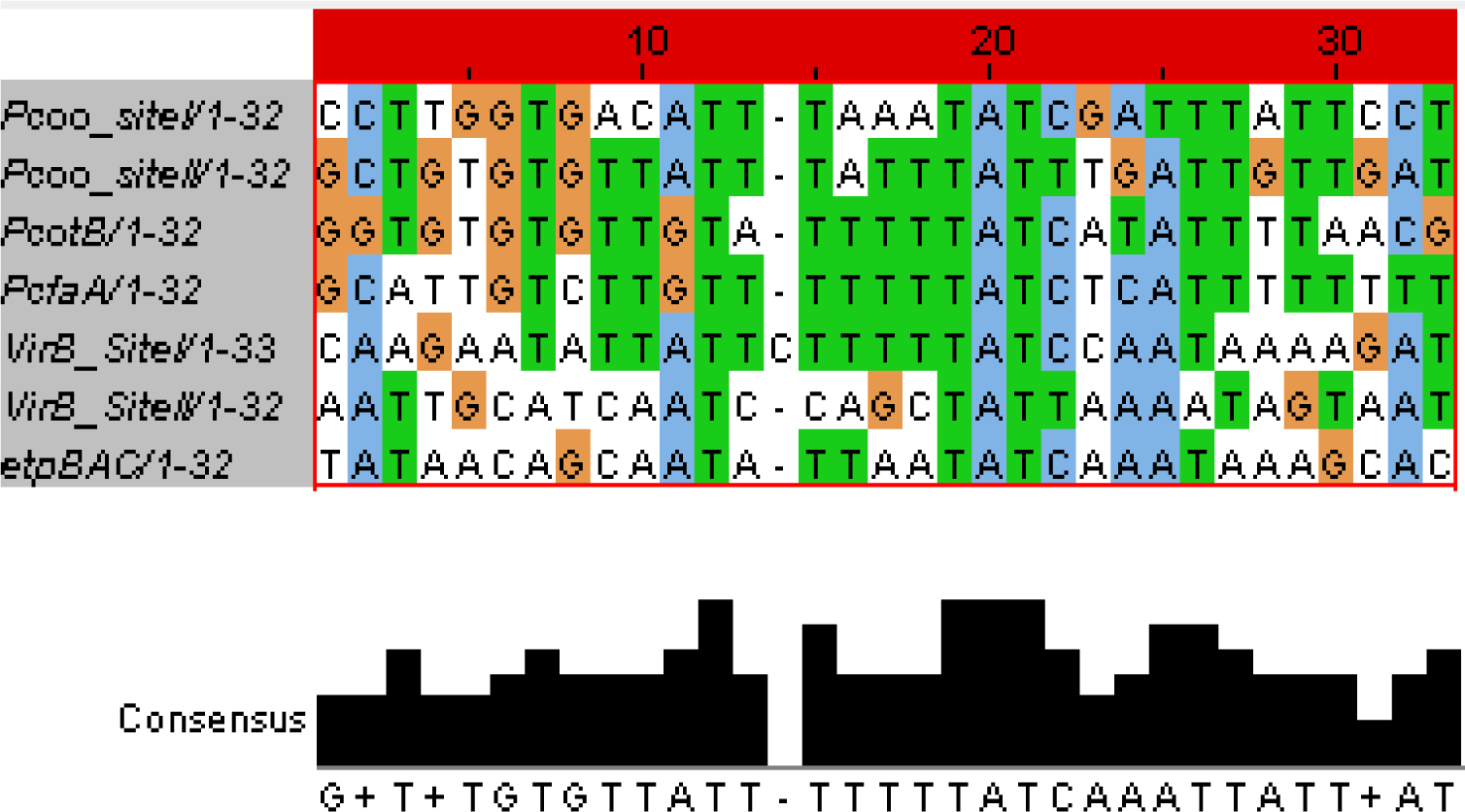
Multiple sequence alignment of Rns binding sites upstream of the *etpBAC* operon and other known binding sites. The Jalview image was generated using Kalign (Lassmann and Sonnhammer, 2005). A consensus sequence is shown at the bottom.

**Fig 3:**
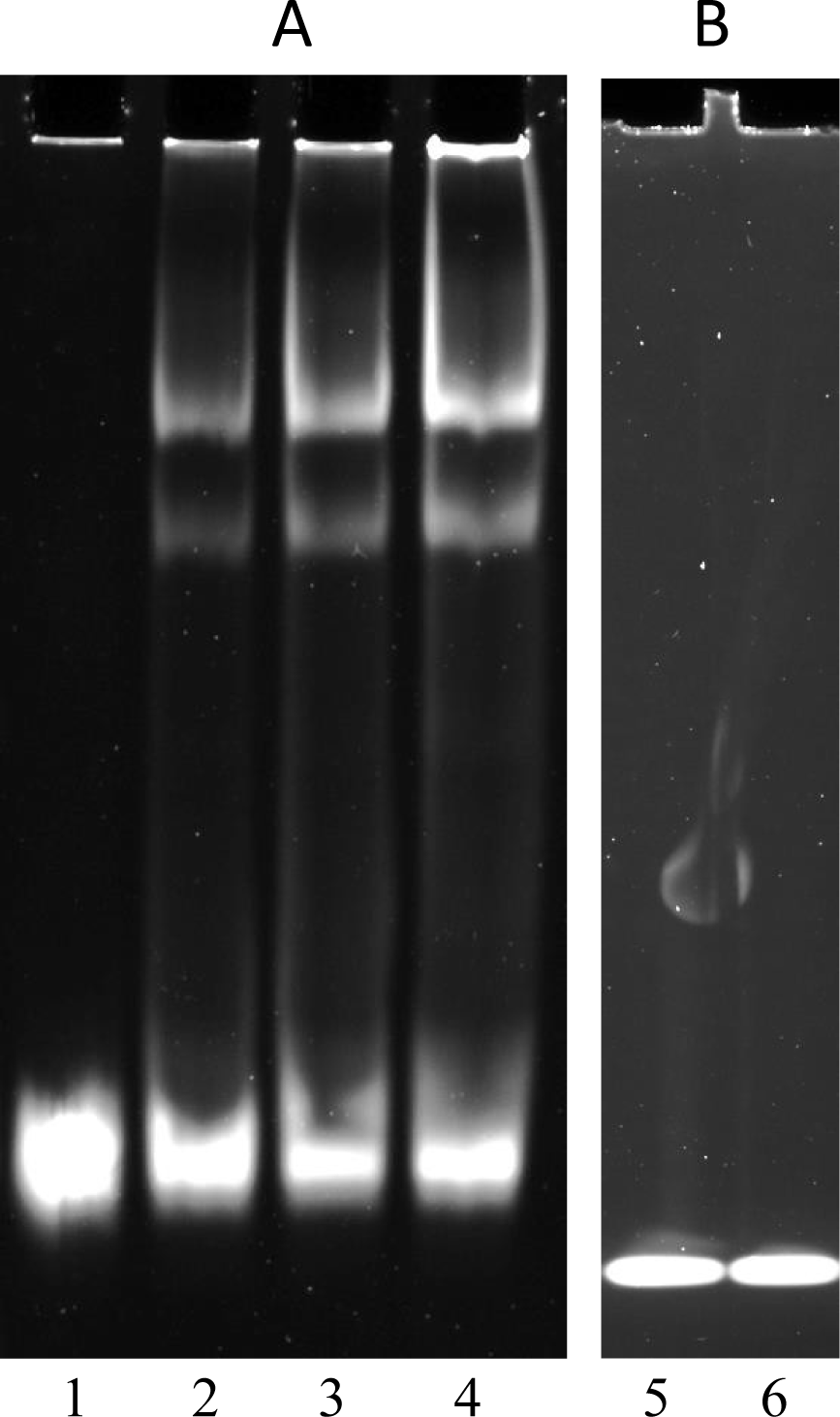
Rns binds to the single predicted site upstream of etpBAC operon. Electrophoretic mobility shift assay (EMSA) of MBP::Rns binding to DNA sequence upstream of *etpBAC* operon (Panel A) or a negative control sequence internal to *gyrB* (Panel B). Lane 1: no MBP::Rns reacted with DNA. Lane 2, 3 and 4: DNA reacted with MBP-Rns at 1 μM, 2 μM and 4 μM, respectively. Lane 5: no MBP-Rns reacted with *gyrA* DNA. Lane 6: *gyrA* DNA reacted with MBP-Rns at a concentration of 4 μM.

## Discussion

Previous studies with AFTRs have revealed their global influence in bacterial gene expression (Hart et al., 2008, Yang et al., 2010). The hypothesis of this project was that Rns exerts a similar influence in the ETEC genome, and influence many of the key virulence factors. In this regard, a combination of different techniques incorporating qRT-PCR and DNA binding assays were employed to investigate this hypothesis.

As an initial step, a single crossover *rns* mutant was created by homologous recombination of a suicide vector into the *rns* locus in B2C Rif^r^. A putative *rns* mutant was confirmed using a PCR analysis for insertional disruption of *rns*, as well as phenotypic and qRT-PCR analysis to confirm the abrogation of *rns* expression. The abolition of haemagglutination and the reduction of *cotB* and *yiiS* transcription were consistent with the insertional disruption of *rns* (data not shown).

As EtpA is known to be a key virulence factors in ETEC, we tested the hypothesis that Rns regulates the expression of *etpB* and *etpA* of *etpBAC*. qRT-PCR analysis of transcription showed that both *etpB* and *etpA* are indeed regulated by Rns. This was a novel finding and is consistent with examples of master regulators of virulence in other bacterial pathogens.

Bioinformatic analysis revealed a potential Rns-binding site sequences upstream of the *etpBAC*. The biological significance of this Rns-binding sequence was confirmed by conducting an EMSA on DNA fragment containing the putative binding. Although the role of the *etpBAC* operon in ETEC virulence (Roy et al., 2009, Meli et al., 2009) has been demonstrated, no regulator of this operon has been elucidated until now. EMSA showed two shifted bands. One possible explanation of this observation is the existence of another binding site in the DNA fragment (Munson et al., 2002). But no such binding site was seen in bioinformatic analysis. A potential alternative explanation could well be found in a recent observation that Rns forms dimers (Mahon et al., 2012). This dual oligomerisation state may lead to two different bands in EMSA. In summary, an important aspect of this work has been the novel finding that Rns positively regulates the expression of non-pilus adhesin, EtpA.

## References

Basturea, G. N., Bodero, M. D., Moreno, M. E. & Munson, G. P. 2008. Residues near the amino terminus of Rns are essential for positive autoregulation and DNA binding. J Bacteriol, 190, 2279–85.

Bodero, M. D., Pilonieta, M. C. & Munson, G. P. 2007. Repression of the inner membrane lipoprotein NlpA by Rns in enterotoxigenic Escherichia coli. J Bacteriol, 189, 1627–32.

Caron, J., Coffield, L. M. & Scott, J. R. 1989. A plasmid-encoded regulatory gene, rns, required for expression of the CS1 and CS2 adhesins of enterotoxigenic Escherichia coli. Proc Natl Acad Sci U S A, 86, 963–7.

Dupont, H. L., Formal, S. B., Hornick, R. B., Snyder, M. J., Libonati, J. P., Sheahan, D. G., Labrec, E. H. & Kalas, J. P. 1971. Pathogenesis of Escherichia coli diarrhea. N Engl J Med, 285, 1–9.

Evans, D. G., Evans, D. J., JR. & Clegg, S. 1980. Detection of enterotoxigenic Escherichia coli colonization factor antigen I in stool specimens by an enzyme-linked immunosorbent assay. J Clin Microbiol, 12, 738–43.

Gallegos, M. T., Schleif, R., Bairoch, A., Hofmann, K. & Ramos, J. L. 1997. Arac/XylS family of transcriptional regulators. Microbiol Mol Biol Rev, 61, 393–410.

Hanahan, D. 1983. Studies on transformation of Escherichia coli with plasmids. J Mol Biol, 166, 557–80.

Hart, E., Yang, J., Tauschek, M., Kelly, M., Wakefield, M. J., Frankel, G., Hartland, E. L. & Robins-Browne, R. M. 2008. RegA, an AraC-like protein, is a global transcriptional regulator that controls virulence gene expression in Citrobacter rodentium. Infect Immun, 76, 5247–56.

Kovach, M. E., Elzer, P. H., Hill, D. S., Robertson, G. T., Farris, M. A., Roop, R. M., 2ND & Peterson, K. M. 1995. Four new derivatives of the broad-host-range cloning vector pBBR1MCS, carrying different antibiotic-resistance cassettes. Gene, 166, 175–6.

Lassmann, T. & Sonnhammer, E. L. 2005. Kalign–an accurate and fast multiple sequence alignment algorithm. BMC Bioinformatics, 6, 298.

Livak, K. J. & Schmittgen, T. D. 2001. Analysis of relative gene expression data using real-time quantitative PCR and the 2(-Delta Delta C(T)) Method. Methods, 25, 402–8.

Mahon, V., Fagan, R. P. & Smith, S. G. 2012. Snap denaturation reveals dimerization by AraC-like protein Rns. Biochimie, 94, 2058–61.

Meli, A. C., Kondratova, M., Molle, V., Coquet, L., Kajava, A. V. & Saint, N. 2009. EtpB is a pore-forming outer membrane protein showing TpsB protein features involved in the two-partner secretion system. J Membr Biol, 230, 143–54.

Munson, G. P., Holcomb, L. G., Alexander, H. L. & Scott, J. R. 2002. In vitro identification of Rns-regulated genes. J Bacteriol, 184, 1196–9.

Munson, G. P. & Scott, J. R. 1999. Binding site recognition by Rns, a virulence regulator in the AraC family. J Bacteriol, 181, 2110–7.

Munson, G. P. & Scott, J. R. 2000. Rns, a virulence regulator within the AraC family, requires binding sites upstream and downstream of its own promoter to function as an activator. Mol Microbiol, 36, 1391–402.

Pilonieta, M. C., Bodero, M. D. & Munson, G. P. 2007. CfaD-dependent expression of a novel extracytoplasmic protein from enterotoxigenic Escherichia coli. J Bacteriol, 189, 5060–7.

Porter, M. E., Mitchell, P., Roe, A. J., Free, A., Smith, D. G. & Gally, D. L. 2004. Direct and indirect transcriptional activation of virulence genes by an AraC-like protein, PerA from enteropathogenic Escherichia coli. Mol Microbiol, 54, 1117–33.

Rice, P., Longden, I. & Bleasby, A. 2000. EMBOSS: the European Molecular Biology Open Software Suite. Trends Genet, 16, 276–7.

Roy, K., Hilliard, G. M., Hamilton, D. J., Luo, J., Ostmann, M. M. & Fleckenstein, J. M. 2009. Enterotoxigenic Escherichia coli EtpA mediates adhesion between flagella and host cells. Nature, 457, 594–8.

Sambrook, J. & Russell, D. W. 2001. Molecular Cloning: a laboratory manual, Cold Spring Harbor, Cold Spring Harbor Laboratory Press.

Yang, J., Tauschek, M., Hart, E., Hartland, E. L. & Robins-Browne, R. M. 2010. Virulence regulation in Citrobacter rodentium: the art of timing. Microb Biotechnol, 3, 259–68.

Yang, J., Tauschek, M. & Robins-Browne, R. M. 2011. Control of bacterial virulence by AraC-like regulators that respond to chemical signals. Trends Microbiol, 19, 128–35.

Zhang, D., Manos, J., Ma, X., Belas, R. & Karaolis, D. K. 2004. Transcriptional analysis and operon structure of the tagA-orf2-orf3-mop-tagD region on the Vibrio pathogenicity island in epidemic V. cholerae. FEMS Microbiol Lett, 235, 199–207.

Zhang, D., Rajanna, C., Sun, W. & Karaolis, D. K. 2003a. Analysis of the Vibrio pathogenicity island-encoded Mop protein suggests a pleiotropic role in the virulence of epidemic Vibrio cholerae. FEMS Microbiol Lett, 225, 311–8.

Zhang, D., Xu, Z., Sun, W. & Karaolis, D. K. 2003b. The vibrio pathogenicity island-encoded mop protein modulates the pathogenesis and reactogenicity of epidemic vibrio cholerae. Infect Immun, 71, 510–5.

